# Brain-scale emergence of slow-wave synchrony and highly responsive asynchronous states based on biologically realistic population models simulated in The Virtual Brain

**DOI:** 10.1101/2020.12.28.424574

**Authors:** Jennifer S. Goldman, Lionel Kusch, Bahar Hazal Yalcinkaya, Damien Depannemaecker, Trang-Anh E. Nghiem, Viktor Jirsa, Alain Destexhe

## Abstract

Understanding the many facets of the organization of brain dynamics at large scales remains largely unexplored. Here, we construct a brain-wide model based on recent progress in biologically-realistic population models obtained using mean-field techniques. We use The Virtual Brain (TVB) as a simulation platform and incorporate mean-field models of networks of Adaptive Exponential (AdEx) integrate-and-fire neurons. Such models can capture the main intrinsic firing properties of central neurons, such as adaptation, and also include the typical kinetics of postsynaptic conductances. We hypothesize that such features are important to a biologically realistic simulation of brain dynamics. The resulting “TVB-AdEx” model is shown here to generate two fundamental dynamical states, asynchronous-irregular (AI) and Up-Down states, which correspond to the asynchronous and synchronized dynamics of wakefulness and slow-wave sleep, respectively. The synchrony of slow waves appear as an emergent property at large scales, and reproduce the very different patterns of functional connectivity found in slow-waves compared to asynchronous states. Next, we simulated experiments with transcranial magnetic stimulation (TMS) during asynchronous and slow-wave states, and show that, like in experimental data, the effect of the stimulation greatly depends on the activity state. During slow waves, the response is strong but remains local, in contrast with asynchronous states, where the response is weaker but propagates across brain areas. To compare more quantitatively with wake and slow-wave sleep states, we compute the perturbational complexity index and show that it matches the value estimated from TMS experiments. We conclude that the TVB-AdEx model replicates some of the properties of synchrony and responsiveness seen in the human brain, and is a promising tool to study spontaneous and evoked large-scale dynamics in the normal, anesthetized or pathological brain.

## 1 Introduction

Conscious and unconscious brain states differ, both at baseline and in response to stimuli, with hallmarks spanning spatio-temporal scales, from neuromodulators acting on molecular membrane channels to changes in global communication between brain regions. Developing a scale integrated understanding of neural dynamics and its effects on computations done by the brain in different states will therefore require linking knowledge spanning ion channel currents (microscale) to the dynamics of brain-wide, distributed, transient functional assemblies (macroscale). Here, using mean-field models of conductance-based, adaptive exponential integrate-and-fire neurons with spike-frequency adaptation developed earlier^1–3^, and constrained by human anatomy and empirically informed local circuit parameters, we report successful simulation of synchronous and asynchronous brain dynamics, thus connecting microscopic to macroscopic scales. Specifically, it has been previously observed that enhanced neuromodulation by neuromodulators such as acetylcholine during active brain states closes K^+^ ion channels, resulting in sustained depolarization of neurons and blocking spike-frequency adaptation (reviewed in ref.^4^). Neuromodulation-induced depolarisation promotes asynchronous, irregular action potential firing. In contrast, low levels of neuromodulators such as acetylcholine during unconscious brain states allow membrane leak channels to open, leading to waves of synchronous depolarisation and hyperpolarisation. In this work, we present a scale integrated model that considers neuromodulation-induced microscopic changes and show that the resulting macroscopic signals are comparable to empirical human data comprising different brain states. This model opens the doors to personalised modelling of human brain states in health and disease, including restful and active waking states, as well as sleep, anaesthesia, and coma.

In this work we aim at simulating full brain activity, during spontaneous activity and after stimulation. Stimulation results in perturbed or evoked activity, with spatiotemporal interactions between areas that have a different fingerprint, corresponding to different brain states. These brain states can be physiological (sleep or awake), pharmacological (e.g. anaesthesia levels), or due to disorders of consciousness (e.g. traumatic brain injury). For these reasons, we have used the simulation capabilities offered by the Human Brain Project’s (HBP’s) EBRAINS neuroscience research infrastructure (https://ebrains.eu and https://ebrains.eu/service/the-virtual-brain) to make access to the models as wide as possible. The simulations illustrate how emergent patterns of activity can be reproduced in silico and shed light on their microscopic underpinnings. These simulations are presented with qualitative and quantitative analyses pioneered in empirical data, for direct comparison to activity recorded during different brain states in actual human subjects^5^.

It is important to note that this work only represents a step in a larger framework, which spans from microscales to macroscales. The first step started with the modeling of biologically-realistic activity states in networks of spiking neurons. Based on experimental recordings, we used the Adaptive Exponential (AdEx) integrate and fire model, which can reproduce two main cell types that can be identified in extracellular recordings, the regular-spiking (RS) and fast-spiking (FS) cells, which can be extracted for example from human recordings^6^. AdEx networks can reproduce *in vivo* activity states^1,7,8^, for asynchronous-irregular (AI) states, as in awake subjects, and Up/Down states as found experimentally in slow-wave sleep^9–11^. An important point for constraining such models was that they interact through realistic biophysical representations of synaptic conductances, which allows the model to be compared to conductance measurements done in awake animals^1^ (for experiments, see^9,12^).

The second step was to obtain mean-field models of AdEx networks interacting with conductances. We used a Master Equation formalism^13^ which was modified to include adaptation^2^. This mean-field model reproduces several features essential to the large scale: (1) the mean-field model captures the level of spontaneous activity, either in AI state (low adaptation) or Up/Down states (high adaptation). This property will be important to simulate different states of spontaneous activity (see Section 3.3). (2) The mean-field model correctly captures the time course of the population response^14^. This feature is important to capture the response of the brain to external stimuli (see Section 3.4). (3) The mean-field model reproduces the fact that the evoked response depends on the activity state of the network^2^. This property of state-dependent responsiveness is central to the work shown here, as we will show that the model can simulate the fact that the response is fundamentally different when the brain is awake (asynchronous mode) or asleep (slow-wave mode)^5^.

Finally, the third step was to test the AdEx mean-field model at mesoscales. We chose to model the millimeter-scale traveling waves found in the primary visual cortex (V1) of awake monkey^15^. A network of mean-field units was constructed to model the V1 traveling waves^1^. The network consisted of several nodes (each represented by an AdEx mean-field) and the connectivity between nodes was adjusted to reproduce the traveling waves occurring following visual stimulation^1^. Importantly, it was also shown that this mesoscale model could capture the properties of the collision between two traveling waves, and in particular the observed suppression which depended on the conductance-based interactions and the different gain of RS and FS cells^16^. Thus, the AdEx mean-field, after being validated at mesoscale, is now ready for being integrated at macroscale, which is the object of the present paper.

## 2 Materials and Methods

We used three types of models, a network of spiking neurons, a mean-field model of this network, and a network of mean-field models implemented in The Virtual Brain (TVB). We describe here these models successively.

### 2.1 Spiking network model

We considered networks of integrate-and-fire neuron models displaying spike-frequency adaptation, based on two previous papers^1,7^. We used the Adaptive Exponential (AdEx) integrate-and-fire model^17^. We considered a population of *N* = 10^4^ neurons randomly connected with a connection probability of *p* = 5%. We considered excitatory and inhibitory neurons, with 20% inhibitory neurons. The AdEx model permits to define two cell types, “regular-spiking” (RS) excitatory cells, displaying spike-frequency adaptation, and “fast spiking” (FS) inhibitory cells, with no adaptation. The dynamics of these neurons is given by the following equations:

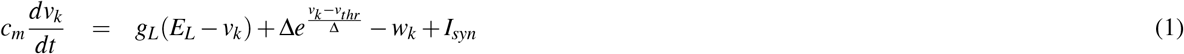

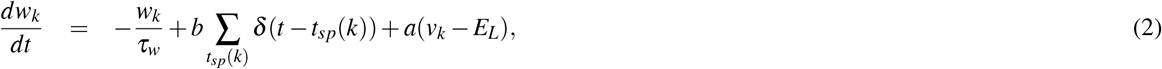

where *c_m_* = 200 pF is the membrane capacitance, *v_k_* is the voltage of neuron *k* and, whenever *v_k_* > *v_thr_* = −50 mV at time *t_sp_*(*k*), *v_k_* is reset to the resting voltage *v_rest_* = −65 mV and fixed to that value for a refractory time *T_refr_* = 5 ms. The leak term *g_L_* had a fixed conductance of *g_L_* = 10 nS and the leakage reversal *E_L_* was of −65 mV. The exponential term had a different strength for RS and FS cells, i.e. Δ = 2mV (Δ = 0.5mV) for excitatory (inhibitory) cells. Inhibitory neurons were modeled as fast spiking FS neurons with no adaptation (*a* = *b* = 0 for all inhibitory neurons) while excitatory regular spiking RS neurons had a lower level of excitability due to the presence of adaptation (while b varied in our simulations we fixed *a* = 4 nS and *τ_w_* = 500 ms if not stated otherwise). The synaptic current *I_syn_* received by neuron *i* is the result of the spiking activity of all neurons *j* ∈ pre(*i*) pre-synaptic to neuron *i*. This current can be decomposed in the synaptic conductances evoked by excitatory E and inhibitory I pre-synaptic spikes

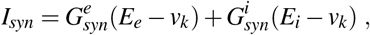

where *E_e_* = 0mV (*E_i_* = −80mV) is the excitatory (inhibitory) reversal potential. Excitatory synaptic conductances were modeled by a decaying exponential function that sharply increases by a fixed amount *Q_E_* at each pre-synaptic spike, i.e.:

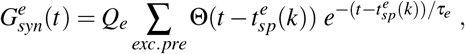

where Θ is the Heaviside function, *τ_e_* = *τ_i_* = 5ms is the characteristic decay time of excitatory and inhibitory synaptic conductances, and *Q_e_* = 1 nS (*Q_i_* = 5 nS) the excitatory (inhibitory) quantal conductance. Inhibitory synaptic conductances are modeled using the same equation with *e* → *i*. This network displays two different states according to the level of adaptation, *b* = 0.005 nA for asynchronous-irregular states, and *b* = 0.02 nA for Up-Down states (see^1^ for details).

### 2.2 Mean-field models

We considered a population model of a network of AdEx neurons, using a Master Equation formalism originally developed for balanced networks of integrate-and-fire neurons^13^. This model was adapted to AdEx networks of RS and FS neurons^1^, and later modified to include adaptation^2^. The latter version is used here, which corresponds to the following equations:

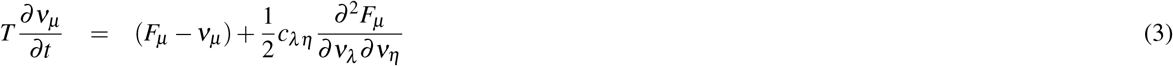

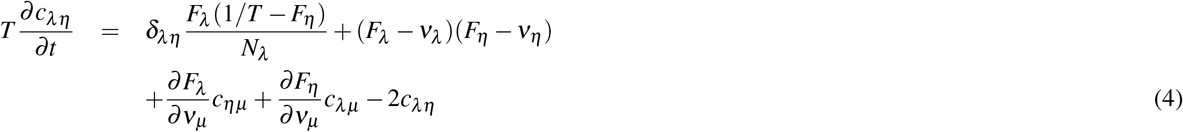

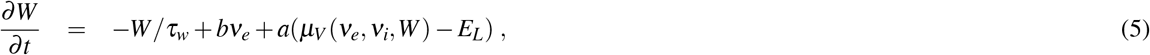

where *μ* = {*e, i*} is the population index (excitatory or inhibitory), *v_μ_* the population firing rate and *c_λη_* the covariance between populations *λ* and *η*. *W* is a population adaptation variable^2^. The function *F*_*μ*={*e,i*}_ = *F*_*μ*={*e,i*}_(*v_e_, v_i_, W*) is the transfer function which describes the firing rate of population *μ* as a function of excitatory and inhibitory inputs (with rates *v_e_* and *v_i_*) and adaptation level *W*. These functions were estimated previously for RS and FS cells and in the presence of adaptation^2^.

At the first order, i.e. neglecting the dynamics of the covariance terms *C_λη_*, this model can be written simply as:

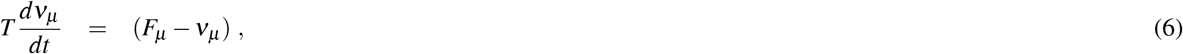

together with Eq. 5. This system is equivalent to the well-known Wilson-Cowan model^18^, with the specificity that the functions *F* need to be obtained according to the specific single neuron model under consideration. These functions were obtained previously for AdEx models of RS and FS cells^1,2^ and the same are used here.

For a cortical volume modeled as a two populations of excitatory and inhibitory neurons, the equations can be written as:

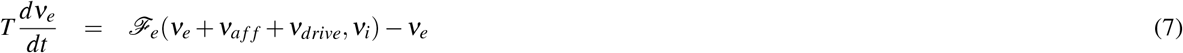

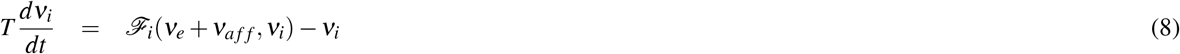

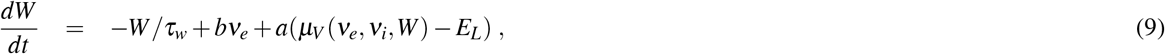

where *ν_aff_* is the afferent thalamic input to the population of excitatory and inhibitory neurons and *ν_drive_* is an external noisy drive. The function *μ_V_* is the average membrane potential of the population and is given by

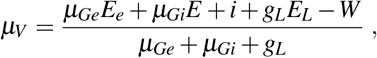

where the mean excitatory conductance is *μ_Ge_* = *ν_e_K_e_τ_e_Q_e_* and similarly for inhibition.

This system describes the population dynamics of a single isolated cortical column, and was shown to closely match the dynamics of the spiking network^2^.

### 2.3 Networks of mean-field models

Extending our previous work at the mesoscale^2,16^ to model large brain regions, we define networks of mean-field models, representing interconnected cortical columns (each described by a mean-field model). For simplicity, we considered only excitatory interactions between cortical columns, while inhibitory connections remain local to each column. The equations of such a network, expanding the two-population mean-field (Eq. 7), are given by:

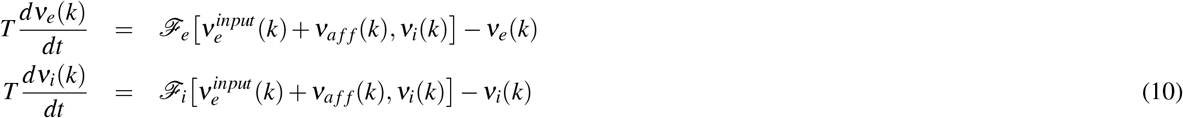

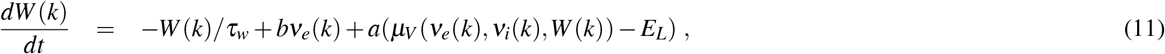

where *ν_e_*(*k*) and *ν_i_*(*k*) are the excitatory and inhibitory population firing rates at site *k*, respectively, *W*(*k*) the level of adapation of the population, and 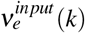 is the excitatory synaptic input. The latter is given by:

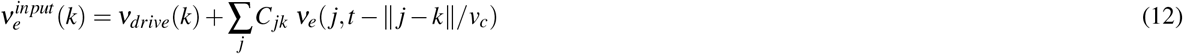

where the sum runs over all nodes *j* sending excitatory connections to node *k*, and *C_jk_* is the strength of the connection from *j* to *k* (and is equal to 1 for *j* = *k*). Note that *ν_e_*(*j, t* – ║*j – k*║ /*v_c_*) is the activity of the excitatory population at node *k* at time *t* – ║*j – k*║/*v_c_* to account for the delay of axonal propagation. Here, ║*j – k*║ is the distance between nodes *j* and *k* and *v_c_* is the axonal propagation speed.

## 3 Results

We start by showing the essential properties of the components forming the TVB-AdEx model. Next, we show the spontaneous dynamics of the TVB-AdEx model, and in the last section we examine the responses to external input.

### 3.1 Components of the TVB-AdEx model

The first component of the TVB-AdEx model is the network of spiking neurons. As shown in previous studies^1,2,7^, networks of AdEx neurons with adaptation can display AI states or Up-Down states, if they are endowed with additional noise (here called “drive”; see Materials and Methods). Figure 1A shows an example of such AI state and Up-Down state dynamics simulated by the same AdEx network for different levels of spike-frequency adaptation (parameter *b* in the equations). In these dynamics the firing of individual units remain very irregular, and of relatively low frequency (Fig. 1B; range of 1-5 Hz for RS cells, 5-20 Hz for FS cells). Note that these firing rates remain very low compared to the maximal frequency allowed by the refractory period (200 Hz in this model).

**Figure 1.**
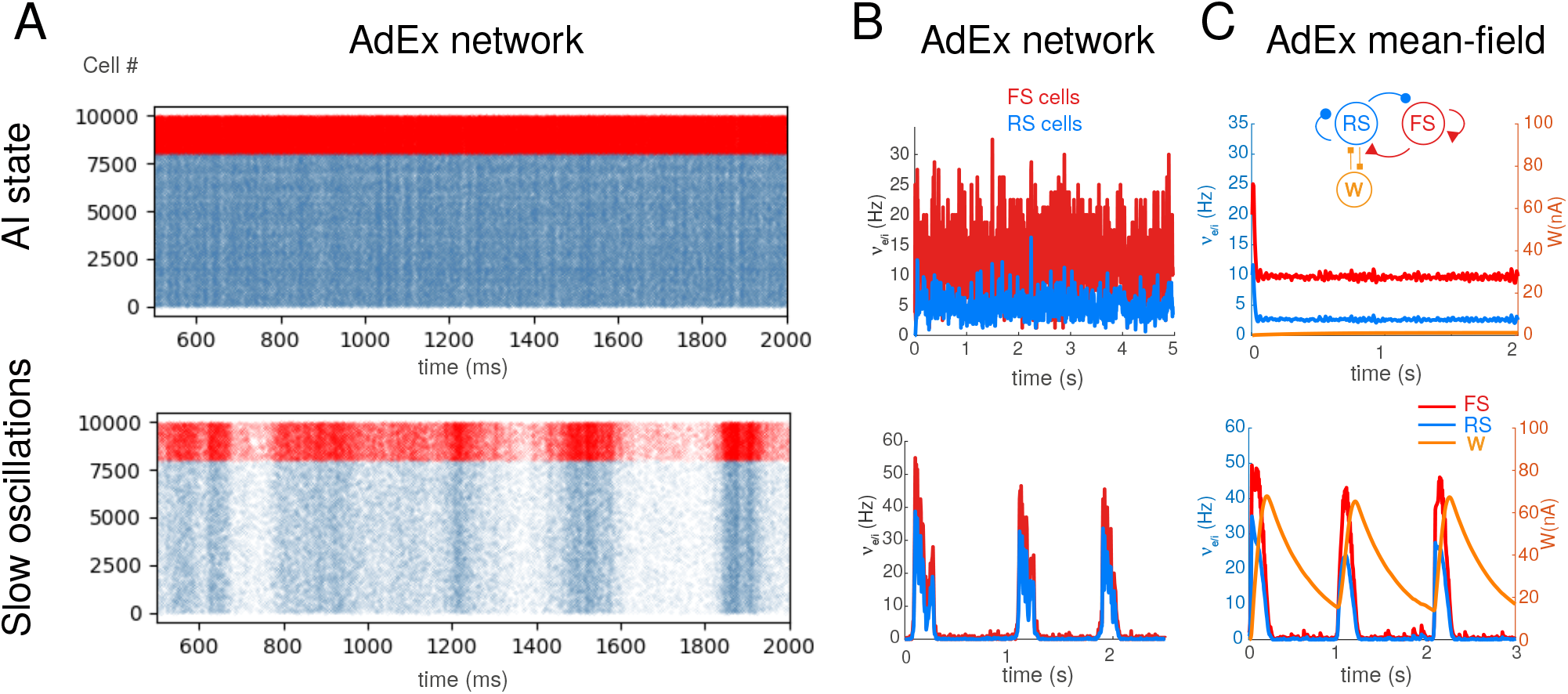
Asynchronous-irregular and slow oscillatory states in AdEx networks and mean-field mdoels. A. Raster plots of excitatory RS (blue) and inhibitory FS (red) AdEx neurons in a network during asynchronous-irregular (AI) state (top), and slow oscillations with Up and Down states (bottom). The two states differed from the valus of adaptation parameter *b* (*b*=1 pA for AI state, and 20 pA for slow oscillations shown here). B. Corresponding mean firing rates of the two populations. C. Mean-field model of AdEx networks with adaptation (scheme in inset). The mean-field reproduces the two states and the corresponding firing rate variations. B-C modified from refs.^2,3^.

The corresponding mean-field model of these two states is shown in Fig. 1C. This model was taken from refs.^2,3^, where it was shown that the mean-field model of AdEx networks with adaptation reproduces these two states, but also their responsiveness^2^. Like the AdEx spiking network, the transition between the two states can be obtained by changing parameter *b*^2^. This mean-field model represents the average dynamics of a cortical column, and will constitute the basis for a single node in the large-scale network.

### 3.2 Integration of mean-field models in TVB

The Virtual Brain^19,20^ is a simulation environment which allows one to integrate the connectivity between different brain regions, estimated for example using diffusion imaging, into a network of many nodes connected according to this connectome. Figure 2 schematizes the integration of the AdEx mean-field models in TVB. We used a human connectome determined from diffusion magnetic resonance imaging collected in the Human Connectome Project, which can be found in https://zenodo.org/record/4263723.X9vvhulKg1J (berlin subjects/DH_20120806). Mean-field models of AdEx networks will be placed at each node of the TVB simulation, and will be connected by excitatory connections according to the connectome. Using such a scheme, is is then possible to simulate large scale networks using AdEx-based mean-field models in TVB, hence the name “TVB-AdEx” model.

**Figure 2.**
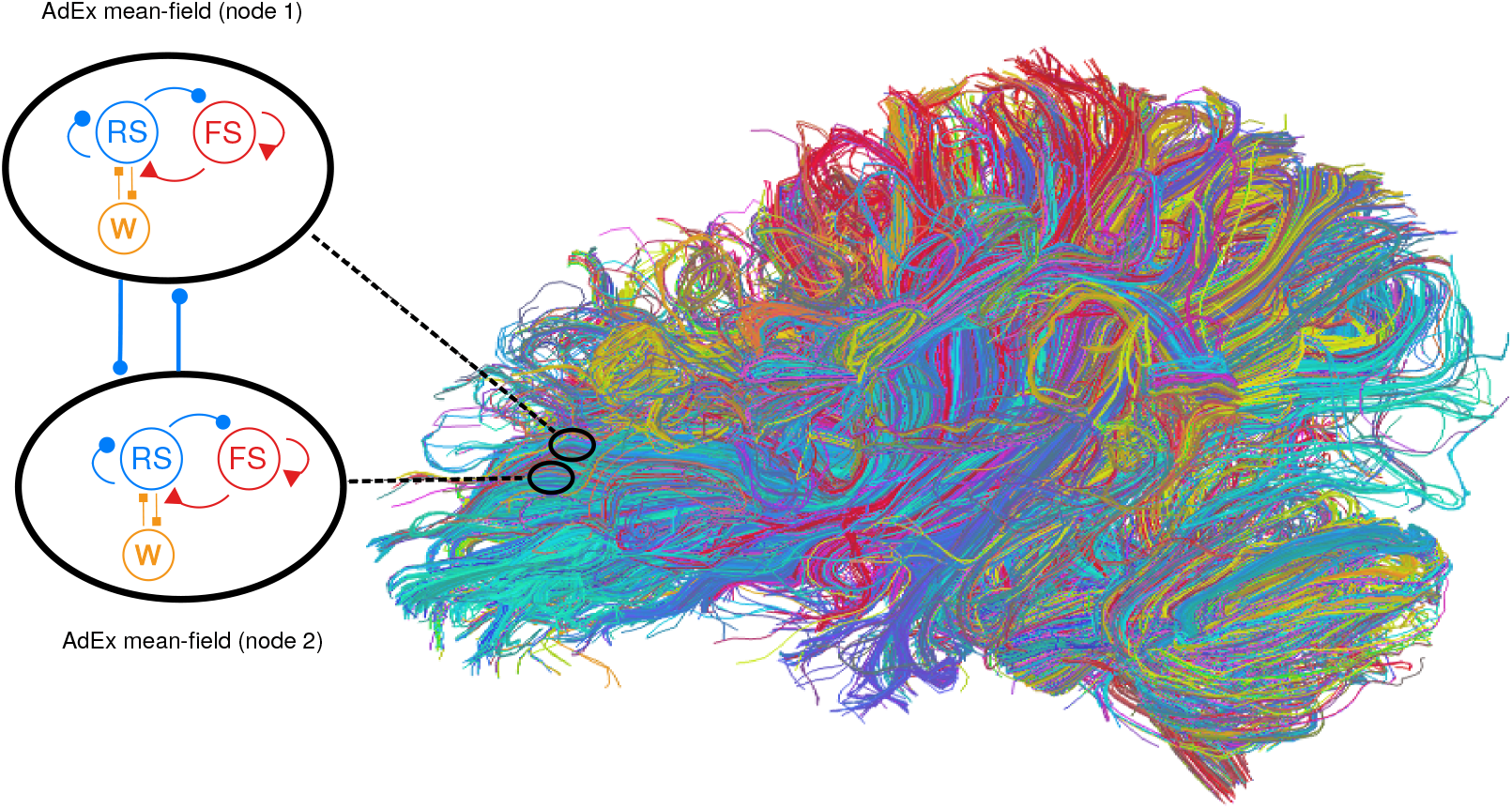
Integration of mean-field models into The Virtual Brain. Left: Each mean-field model consists of two populations of excitatory (blue) and inhibitory (red) neurons, together with an adatation variable (*W*, orange). This mean-field model represents the dynamics of a single node, and different nodes are connected with excitatory connections (thick blue lines). Right: integration of mean-field models in The Virtual Brain (TVB), which can include brain connectivity as defined from diffusion imaging. In this case, a large number of mean-field models are connected according to this connectome.

Once the mean-field models are integrated in each node of the network, the connectivity will depend on the spatial resolution of the coarse-graining. For example, TVB allows simple simulations using a few tens of nodes, as illustrated in Fig. 3. In this case, TVB calculates the connectivity between each node according to the connectome, and the resulting simulation is therefore very coarse-grained, with each mean-field representing a substantially large brain area. TVB can also simulate much finer grained connectivity, by defining a large number of nodes (usually of the order or tens of thousand nodes). Each mean-field then represents a much more local population, depending on the size of the voxel it represents. It makes sense to use voxel size corresponding to fMRI imaging (millimeter scale). The drawback of such fine grained simulations is that they typically require large computing resources, while a coarse grained TVB simulation can be run on a standard workstation.

**Figure 3.**
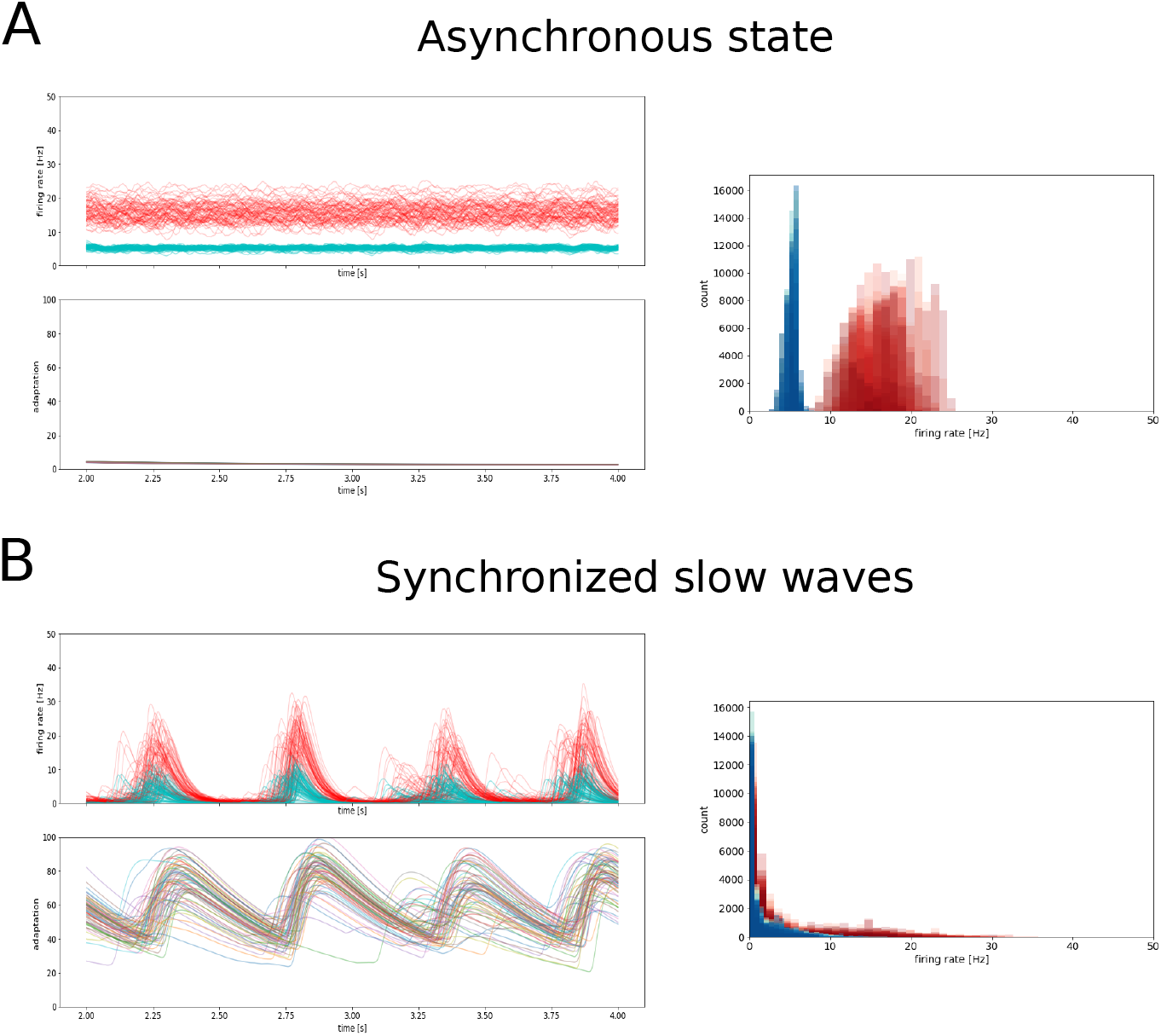
Whole-brain activity during asynchronous and synchronized slow-wave states. A. Asynchronous state. B. Synchronous slow-waves. In each panel, the activity in the different nodes is shown for a few seconds (top left), together with the adaptation (bottom left). The histogram of activities is shown for each case (right panels).

### 3.3 Spontaneous activity of large-scale networks

Figure 3 shows a typical TVB simulation of a 76-node network, where each node was placed in either AI or in Up-Down state conditions. When the mean-field models in each node are set to AI mode (*b*=2 pA), the activity of the whole-brain network sets into an asynchronous mode (Fig. 3A). When they are set into the Up/Down state mode, they tend to synchronize into large-scale slow-waves (Fig. 3B).

The power spectral density of these signals (Fig. 5) shows a different struture in the two states. In the asynchronous state, interestingly, a peak appears around a frequency of 10 Hz (Fig. 5A), an oscillation not present in a single node, and which is thus an emerging property of the network. This oscillation was also observed in the connected mean-fields for sufficiently strong coupling (not shown). In the slow-wave state, a peak appears around 1 Hz (Fig. 5B), in the frequency range of delta waves.

**Figure 4.**
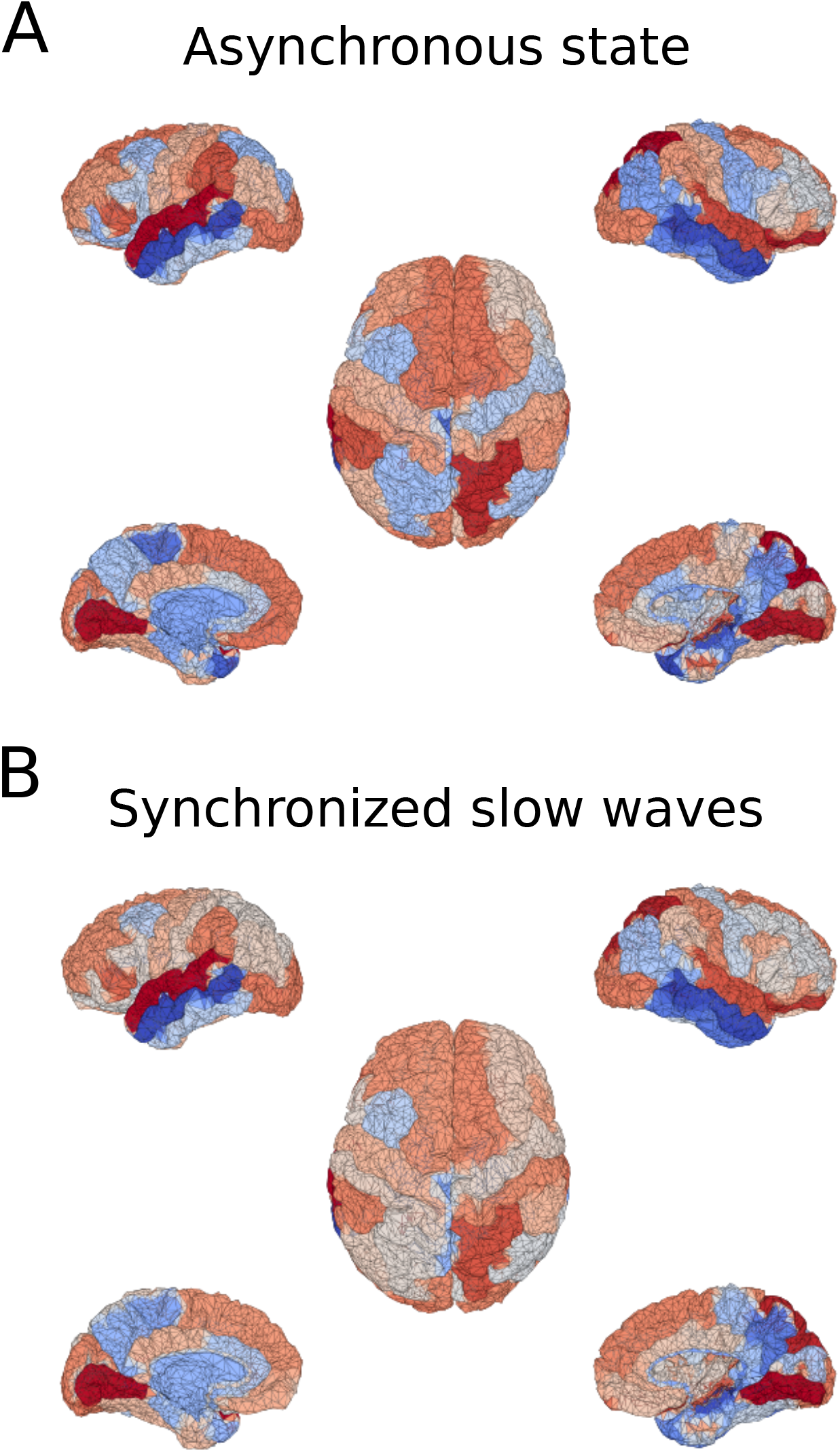
Whole-brain activity maps during asynchronous and synchronized slow-wave states. A. Asynchronous state. B. Synchronous slow-waves. Same data as in Fig. 3 represented as a function of space maps.

**Figure 5.**
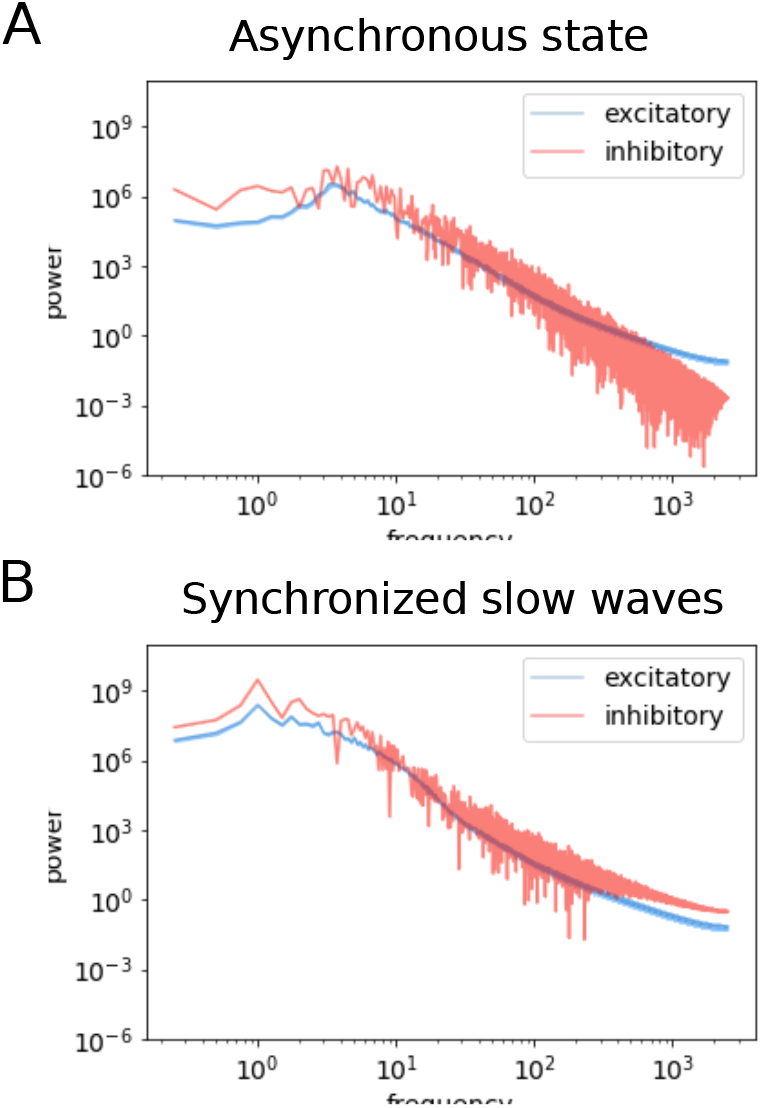
Power spectral density of whole-brain activity during asynchronous and synchronized slow-wave states. A. Asynchronous state. B. Synchronous slow-waves.

To further investigate the spontaneous activity generated by this large-scale network, we computed the correlations and functional connectivity, as measured by the Pearson correlation between mean firing rates in time between pairs of regions for E and I neuron types (Fig. 6). Interestingly, we can see that the correlations remain low when the mean-field model is in the AI state, which defines an asynchronous state at the large-scale network level. In this asynchronous mode, the functional connectivity and structural connectivity are more similar (Fig. 6A; compare with right panel). This is not the case, however, in Up-Down states, where the large-scale network tends to synchronize the Up and Down states generated by the local mean-field models. This synchrony is a typical feature of sleep slow-waves^10,11^. This corroborates the synchrony that is already visible in the activity (see Fig. 3), which is an emergent property of the large-scale network when the individual nodes are set to the slow-wave mode. In this case, the structure of functional connectivity is different from the structure of physical connectivity (Fig. 6B).

**Figure 6.**
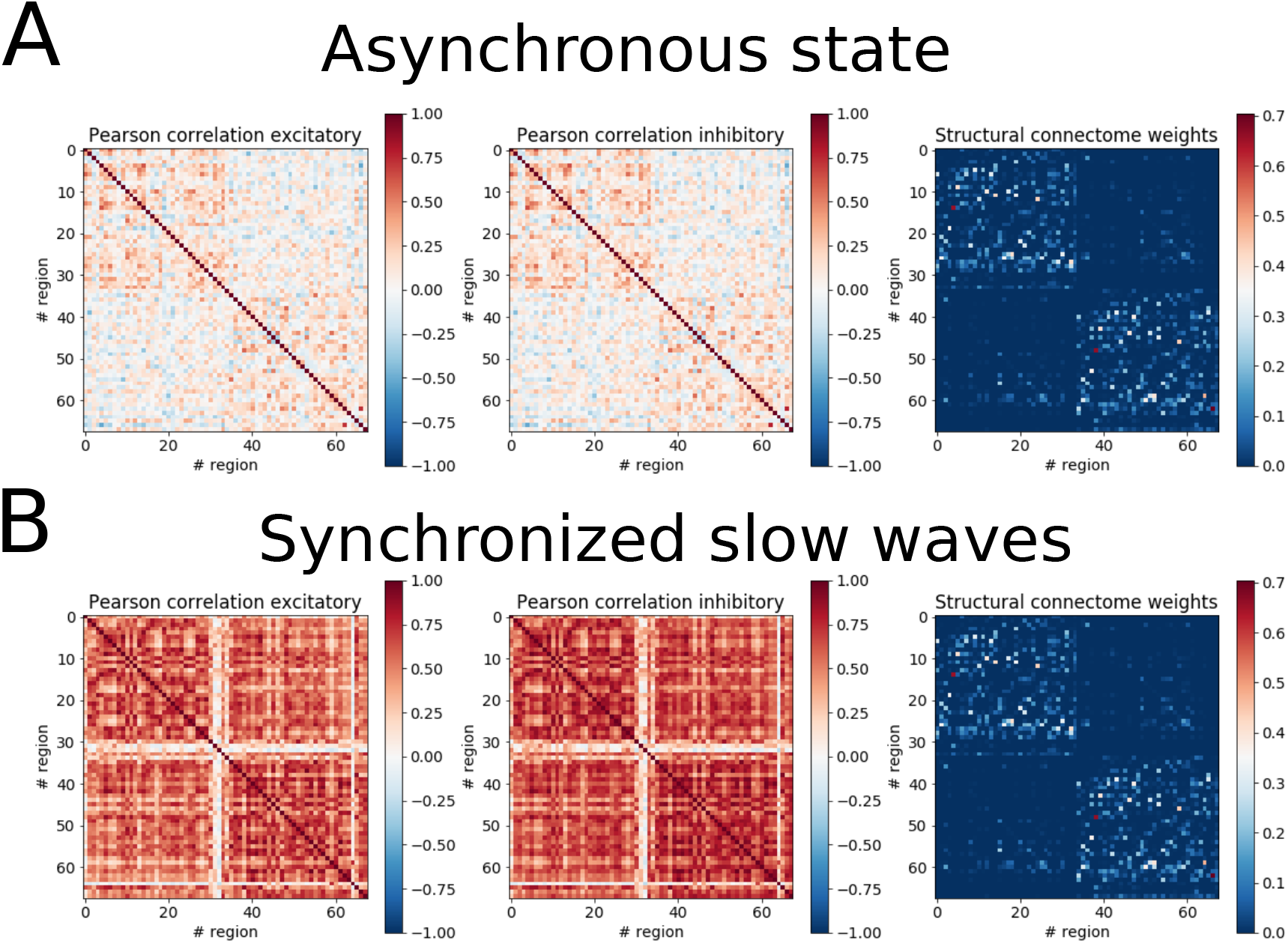
Functional connectivity in synchronized and desynchronized states. A. Asynchronous state, B. Synchronized slow-waves. The pearson correlation between brain regions is shown for excitatory activities (left) and between inhibitory populations (middle). The structural connection weights (extracted from the connectome) are shown for comparison (right).

**Figure 7.**
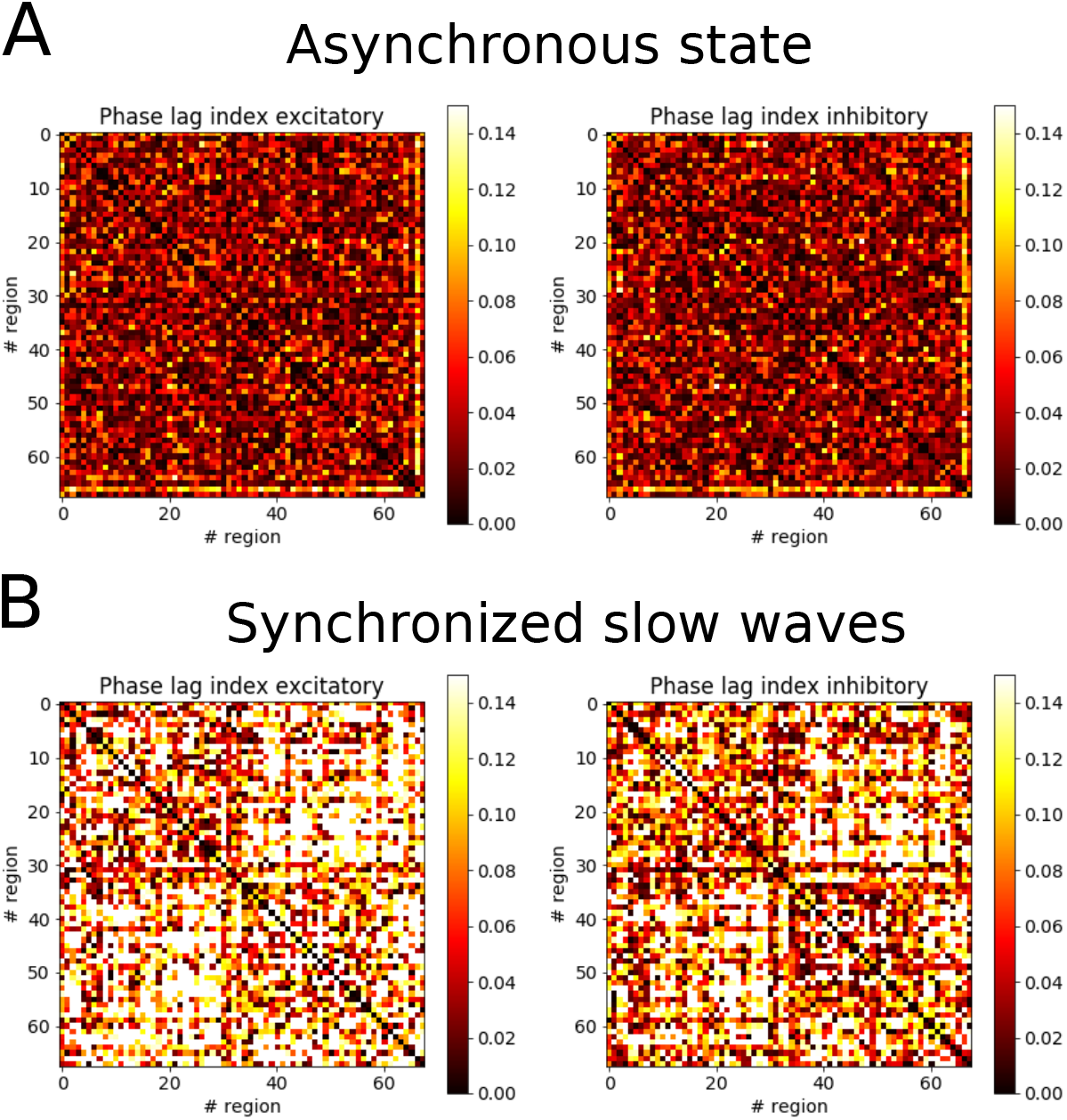
Phase-locking index (PLI) in synchronized and desynchronized states. A. Asynchronous state, B. Synchronized slow-waves. The PLI was calculated between the mean firing rate activities between different regions, represented similarly as in Fig. 6, for excitatory activities (left) and between inhibitory populations (right).

### 3.4 Response to external stimulation

Figure 8 illustrates the effect of external stimulation in the large-scale network defined by the TVB-AdEx model. The effect of the external stimulus is apparent for both Up-Down states (Fig. 8A) and the asynchronous state (Fig. 8B). The average responses (Fig. 9) show that the response is typically of high amplitude in Up-Down states, while it is of lower amplitude in asynchronous states. This feature corresponds to experimental observations^5^.

**Figure 8.**
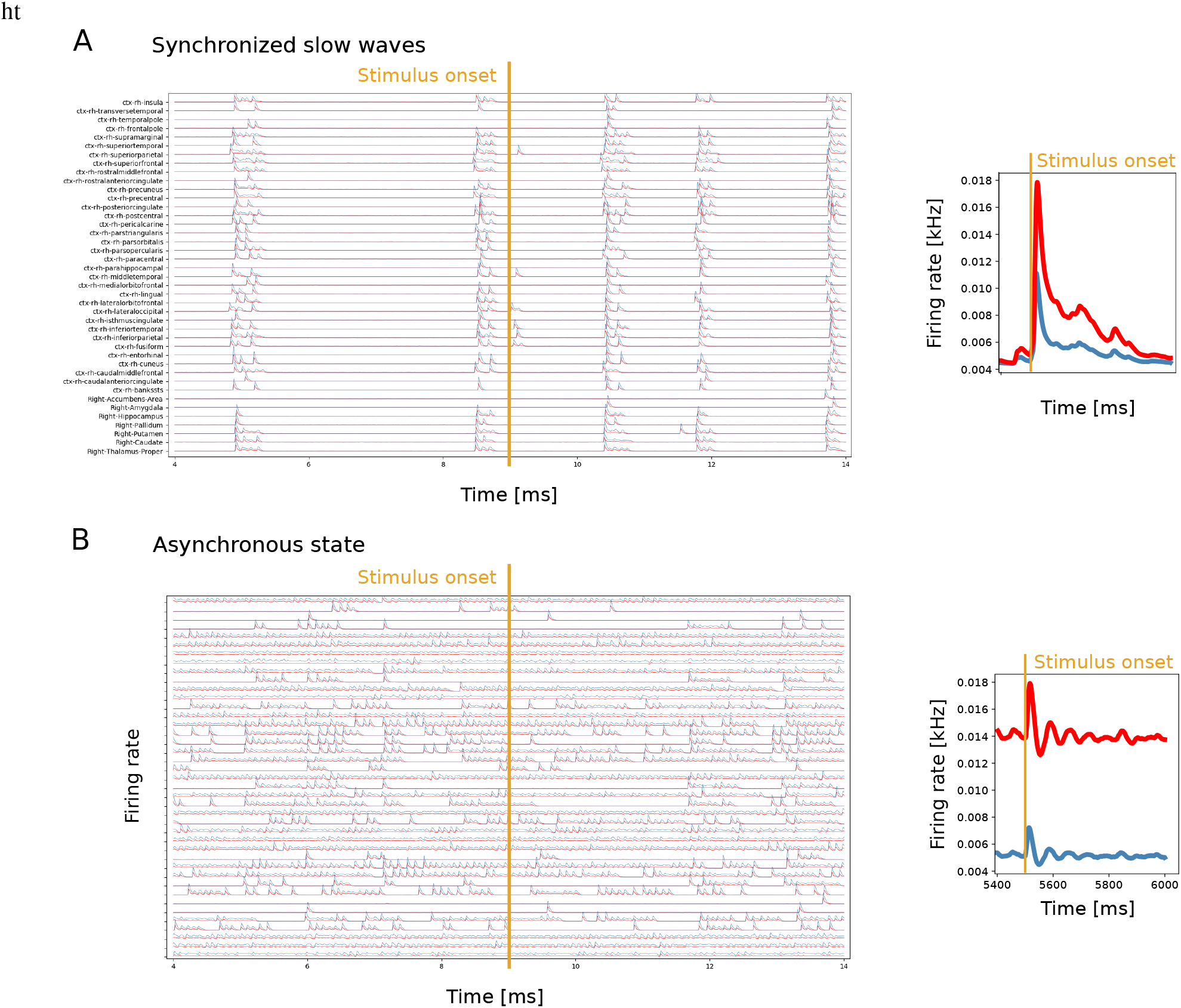
Response of the TVB-AdEx model to external stimulation. The same stimulation is given in Up-Down states (A) or in the asynchronous state (B). The left panel shows selected brain regions during one example stimulation. The right panel displays the average response to 100 stimuli. The stimulus was given here in the occipital cortex.

**Figure 9.**
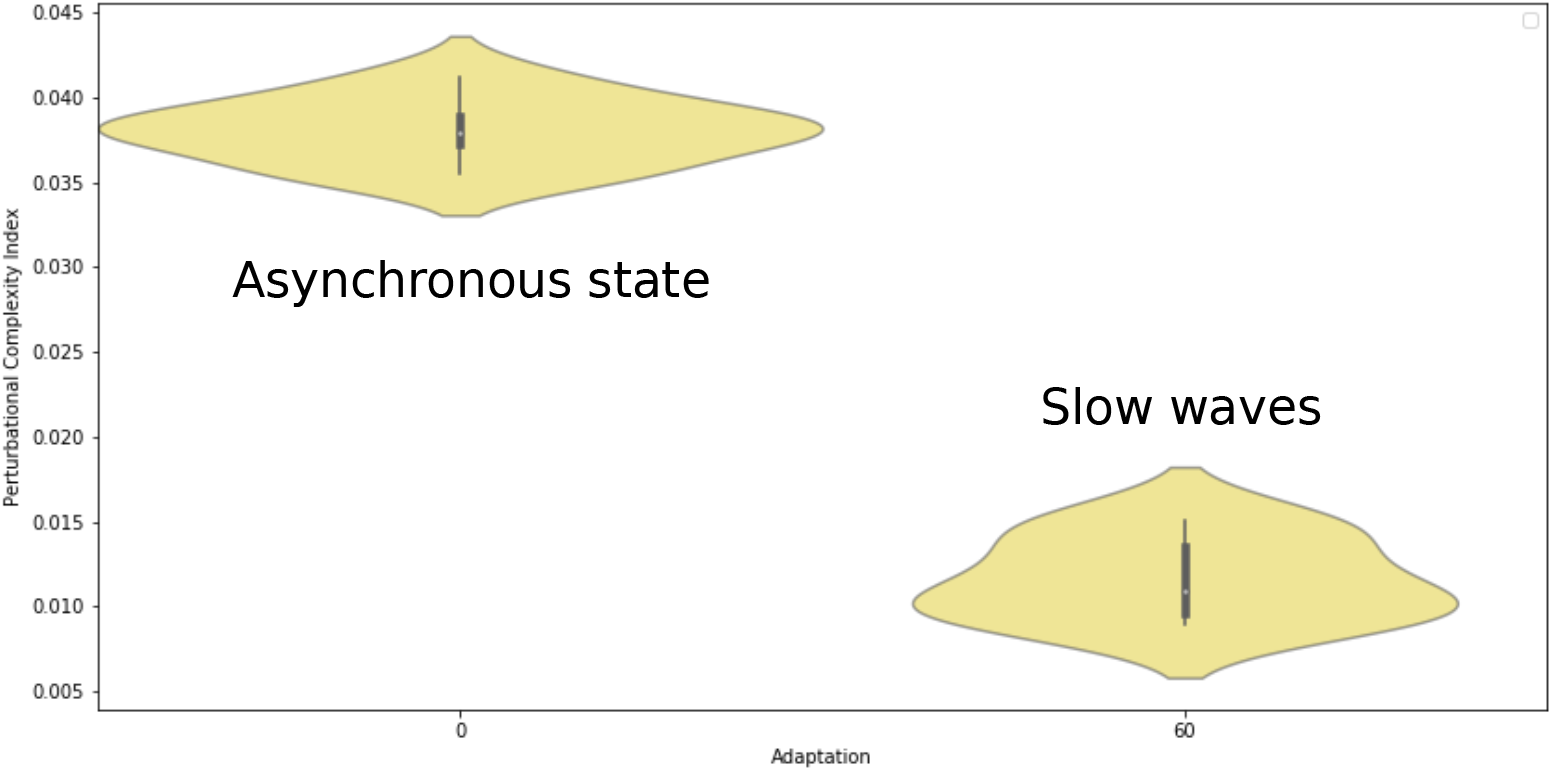
Perturbational complexity index (PCI) in two different simulated brain states. The PCI was high during the asynchronous state, and was low during slow-waves.

To better characterize the spatiotemporal effect of the stimulus, we have computed the same measure as in the experiments, the perturbational complexity index (PCI). A low PCI value indicates a “simple” response, while a high PCI value indicates more “complex” responses, typically propagating to different brain areas^5^. The propagation of activity in different regions of the brain is illustrated in Fig. 10.

**Figure 10.**
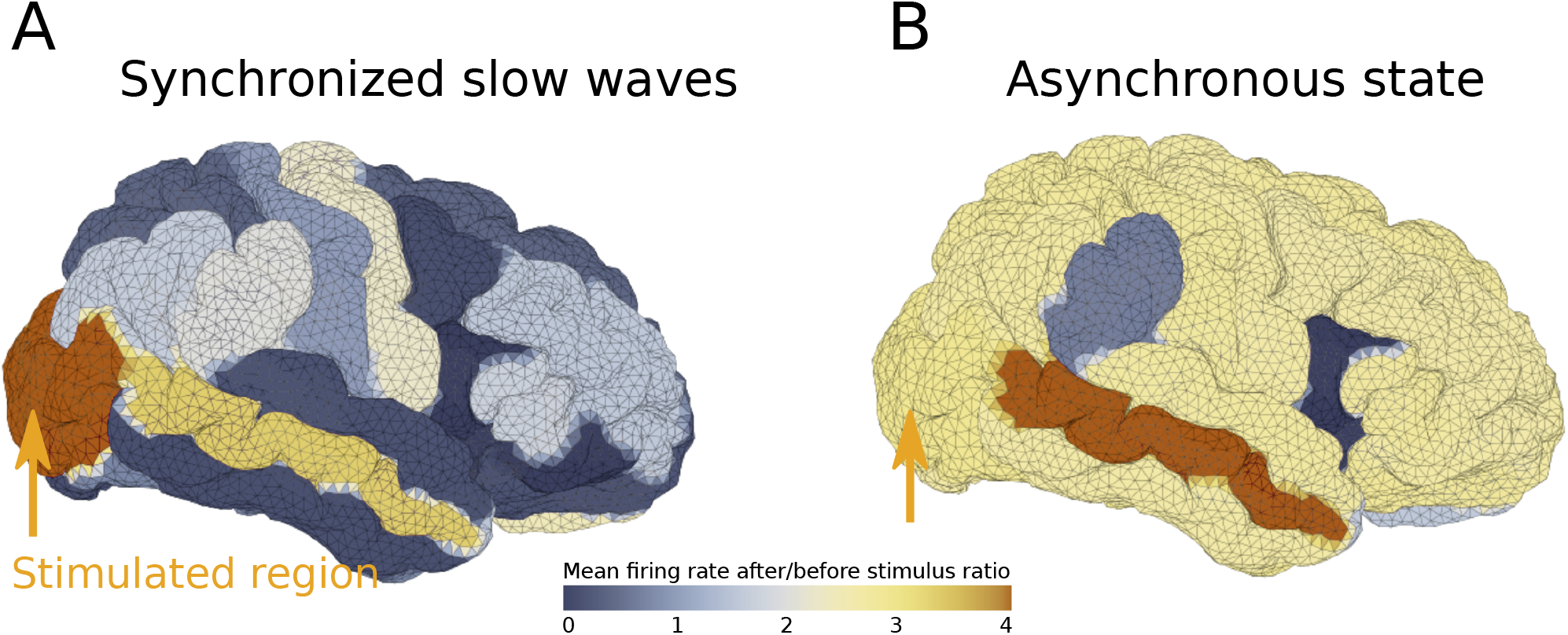
Propagation of evoked activity in the brain simulated with the TVB-AdEx model. The ratio of activity after and before the stimulus was computed and represented here according to a color scale across the simulated brain, for Up-Down state dynamics (A) and the asynchronous state (B).

This is what we observe in the TVB-AdEx model. As shown in Fig. 9, computing the PCI gives values that are typically high for asynchronous states, and lower in more synchronized states displaying Up-Down activity. The same behavior was observed when comparing awake subjects with subjects in slow-wave sleep^5^. The different spread of the activity in the two states is also illustrated in Fig. 10.

## 4 Discussion

In this paper, we have shown the use of “biologically-realistic” mean-field models^2^ to simulate large-scale networks using the TVB platform. The mean-field models were derived from networks of AdEx neurons displaying asynchronous or Up-Down states. After integration in TVB, the resulting TVB-AdEx model displays a number of interesting features, and several exciting perspectives for future work, which we discuss here.

A first result is that we were able to identify “synchronized” and “desynchronized” states based on this integration. The TVB-AdEx model, when set into the Up-Down state (with high adaptation), displays synchronized Up and Down states at a large network level (Fig. 3). This is consistent with the synchronized Up-Down state dynamics observed during slow-wave activity in the brain^10,11,21^. When the same model is set into the asychronous-irregular regime, the large-scale network displays a much lower level of synchrony (Fig. 3), consistent with the asynchronous dynamics typically seen in awake and aroused states^10,11,21^. These are emergent properties of the large-scale network.

A second main result is that the evoked dynamics is also state dependent, in a manner consistent with the experimental data. External stimuli of different amplitudes propagate differently according to the state of the network. When the network displays synchronized Up and Down states, the stimulus typically evokes a high-amplitude response, but that remains local to a close neighborhood of the stimulation site. When it is in the asynchronous regime, the same stimuli evoke responses that are weaker in amplitude, but that propagate in a much more elaborated way. The PCI measure applied to these two states match the experimental observations^5^. Here again, this is an emerging property of the large-scale network.

What are possible mechanisms for such differences? A previous study^8^ showed that not all states are equal in balanced networks, and that asynchronous states, despite their apparently noisy character, can display higher responsiveness and support propagation across layers. This higher responsiveness of AI states can be explained by the combined effects of depolarization, membrane potential fluctuations and conductance state. It was proposed as a fundamental property to explain why the activity of the brain is systematically asynchronous in aroused states^8^. The present results are in full agreement with this mechanism, which manifests here in the asynchronous state as a propagation across many brain areas, which is associated with high values of the PCI.

We believe that this work opens several perspectives. First, the enhanced propagation of stimuli during asynchronous states could be used as a basis to explain why stimuli are perceived in asynchronous states, and what kind of modulation of the network activity could support phenomena such as perception or attention. Second, mean-field models can be set to also display pathological states, such as hyperactive or hypersynchronized states, and the TVB-AdEx model could be used to investigate seizure activity. Other features could also be included in mean-field models, such as neuronal heterogeneity^14^, and further increase their biological realism.

Finally, the anatomical backbone and functional parameters of this model can be substituted to study other animals. For example, the Allen Institute offers a rich database of neuronal connectivity from axonal tracing studies in mouse (https://alleninstitute.org), providing a ground truth to anchor connectivity determined in neuroimaging experiments, as well as providing models adapted to better understanding experiments performed in mice. The present approach of a whole-brain model based on biophysically-informed properties summarising microscopic to macroscopic dynamics and their differences between brain states could be applied to the mouse brain and to any other species for which the whole-brain white-matter connectivity is available.

## Acknowledgments

This work was supported by the Centre National de la Recherche Scientifique (CNRS, France), the European Community Future and Emerging Technologies program (Human Brain Project, H2020-785907), the ANR PARADOX, and the ICODE excellence network.

## Code availability

A python-based open-access code to run the present whole-brain model will be accessible online in the EBRAINS platform (https://ebrains.eu) as a companion to the publication of the present article.

